# New insights from Thailand into the maternal genetic history of Mainland Southeast Asia

**DOI:** 10.1101/162610

**Authors:** Wibhu Kutanan, Jatupol Kampuansai, Andrea Brunelli, Silvia Ghirotto, Pittayawat Pittayaporn, Sukhum Ruangchai, Roland Schröder, Enrico Macholdt, Metawee Srikummool, Daoroong Kangwanpong, Alexander Hübner, Leonardo Arias Alvis, Mark Stoneking

**Affiliations:** Department of Biology, Faculty of Science, Khon Kaen University, Khon Kaen, Thailand; Department of Evolutionary Genetics, Max Planck Institute for Evolutionary Anthropology, Leipzig, Germany; Department of Biology, Faculty of Science, Chiang Mai University, Chiang Mai, Thailand; Department of Life Science and Biotechnology, University of Ferrara, Ferrara, Italy.; Department of Linguistics, Faculty of Arts, Chulalongkorn University, Bangkok, Thailand; Material Science and Nanotechnology Program, Faculty of Science, Khon Kaen University, Khon Kaen, Thailand; Department of Biochemistry, Faculty of Medical Science, Naresuan University, Phitsanulok, Thailand

**Keywords:** mitochondrial genome, central Thai people, demic diffusion, Tai-Kadai, Austronesian

## Abstract

Tai-Kadai (TK) is one of the major language families in Mainland Southeast Asia (MSEA), with a concentration in the area of Thailand and Laos. Our previous study of 1,234 mtDNA genome sequences supported a demic diffusion scenario in the spread of TK languages from southern China to Laos as well as northern and northeastern Thailand. Here we add an additional 560 mtDNA sequences from 22 groups, with a focus on the TK-speaking central Thai people and the Sino-Tibetan speaking Karen. We find extensive diversity, including 62 haplogroups not reported previously from this region. Demic diffusion is still a preferable scenario for central Thais, emphasizing the extension and expansion of TK people through MSEA, although there is also some support for an admixture model. We also tested competing models concerning the genetic relationships of groups from the major MSEA languages, and found support for an ancestral relationship of TK and Austronesian-speaking groups.

## Introduction

The geography of Thailand encompasses both upland and lowland areas, and Thailand is one of the most ethnolinguistically-diverse countries in Mainland Southeast Asia (MSEA). With a census size of ~ 68 million in 2015, there are 70 different recognized languages belonging to five different major language families: Tai-Kadai (TK) (90.5%), Austroasiatic (AA) (4.0%), Sino-Tibetan (ST) (3.2%), Austronesian (AN) (2.0%), and Hmong-Mien (HM) (0.3%) (Simons and Fennig, 2017). The majority of the people (29.72%) are called Thai or Siamese and speak a central Thai (CT) language that belongs to the TK family. Since it is the country’ s official language, the number of people speaking the CT language as their primary or secondary language is ~ 40 million (Simons and Fennig, 2017), or ~ 68% of the population.

The recorded history of the CT people or Siamese started with the Sukhothai Kingdom, around the 13^th^ century A.D. (Baker and Phongpaichit, 2009). However, before the rise of the TK civilization, Thailand was under the control of Mon and Khmer people (Revire, 2014; Baker and Phongpaichit, 2017). Linguistic and archaeological evidence suggests that the prehistorical TK homeland was situated in the area of southeastern or southern China, and that they then spread southward to MSEA around 1-2 kya (O’ Connor, 1995; Pittayaporn, 2014). This process could have occurred via demic diffusion (i.e., a migration of people from southern China, who are then the ancestors of present-day CT people), cultural diffusion (i.e., the CT ancestors were AA groups who shifted to TK languages), or admixture (i.e., a migration of people from southern China who admixed with AA groups, so CT people have ancestry from both sources). We previously used demographic modeling to test these scenarios, using a large dataset of complete mtDNA genome sequences from Thai/Lao people, mostly from northern and northeastern Thailand, and found support for the demic diffusion model (Kutanan et al., 2017). However, CT groups were not included in that study, and could have a different history.

Here we extend our previous study by adding 560 new complete mtDNA genome sequences from 22 groups (mostly from CT) speaking TK, AA, and ST languages; when combined with the previous data (Kutanan et al. 2017), there are a total of 1,794 sequences from 73 Thai/Lao groups. We find extensive diversity in the new groups, including 62 haplogroups not found in the previous study. We use demographic modeling to test three competing scenarios (demic diffusion, cultural diffusion, and admixture) for the origins of CT groups. We also use demographic modeling to test competing scenarios (Peiros, 1998; Sagart, 2004; 2005; Starosta, 2005) for the genetic relationships of groups speaking languages from the major MSEA language families (TK, AA, ST and AN). Our results provide new insights into the maternal genetic history of MSEA populations.

## Materials and Methods

### Samples

Samples were analyzed from 560 individuals belonging to 22 populations classified into four groups: 1) the central Thais (7 populations: CT1-CT7); 2) the Mon (2 populations: MO6-MO7); 3) the TK speaking groups from northern Thailand, including Yuan (4 populations: YU3-YU6), Lue (4 populations: LU1-LU4) and Khuen (TKH); and 4) the ST speaking Karen (4 populations: KSK1, KSK2, KPW and KPA) (Table 1 and Figure 1). Genomic DNA samples of MO6, Yuan, Lue, Khuen and Karen were from previous studies (Kampuansai et al., 2007; Lithanatudom et al., 2016) while the MO7 and central Thai groups were newly-collected saliva samples obtained with written informed consent. DNA was extracted by QIAamp DNA Midi Kit(Qiagen, Germany). This research was approved by Khon Kaen University, Chiang Mai University, Naruesuan University, and the Ethics Commission of the University of Leipzig Medical Faculty.

**Table 1.**
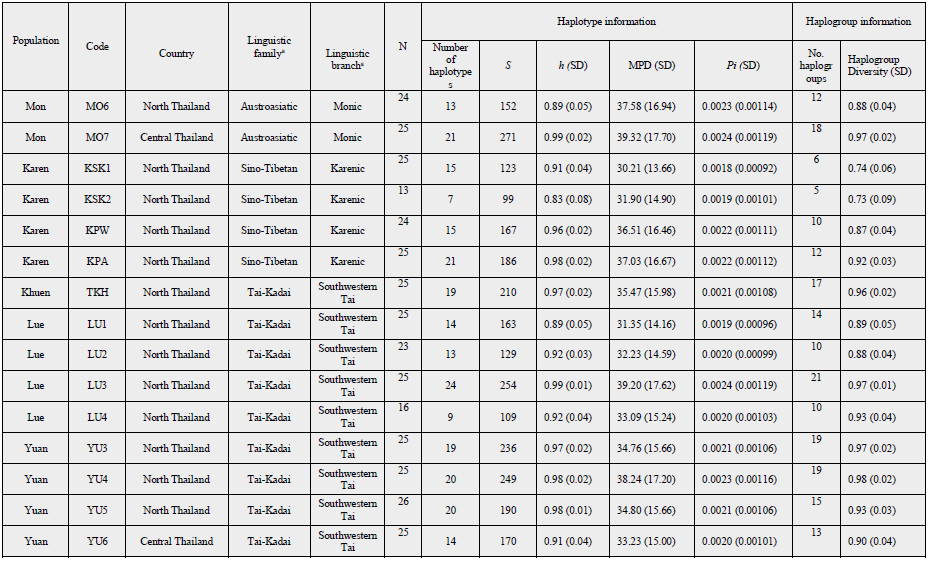
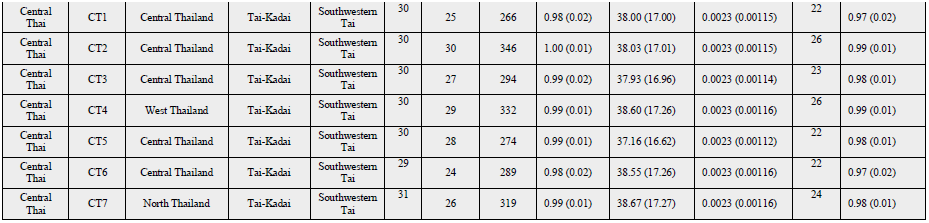
Population information and summary statistics

**Figure 1.**
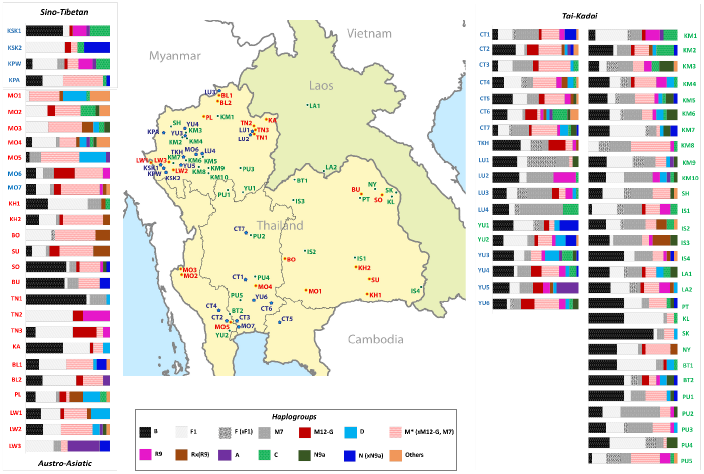
Map showing sample locations and haplogroup distributions. Blue stars indicate the 22 presently studied populations (Tai-Kadai, Austroasiatic and Sino-Tibetan groups) while red and green circles represent Tai-Kadai and Austroasiatic populations from the previous study (Kutanan et al., 2017). Population abbreviations are in Supplementary Table 1.

### Sequencing

We generated complete mtDNA sequences from genomic libraries with double indices and mtDNA enrichment based on protocols described previously (Meyer and Kircher, 2010; Maricic et al., 2010). The libraries were sequenced on the Illumina Hiseq 2500. MtDNA consensus sequences were obtained as described by Arias-Alvis et al. (2017) except that Illumina standard base calling was performed using Bustard and the read length was 76 bp. Sequences were manually checked with Bioedit (<u>www.mbio.ncsu.edu/BioEdit/bioedit.html</u>). A multiple sequence alignment of the sequences and the Reconstructed Sapiens Reference Sequence (RSRS) (Behar et al., 2012) was obtained by MAFFT 7.271 (Katoh and Standley, 2013).

### Statistical Analyses

Haplogroup assignment was performed with the online tools Haplogrep (Kloss-Brandstä tter et al., 2011) and MitoTool (Fan and Yao, 2011). Arlequin 3.5.1.3 (Excoffier and Lischer, 2010) was used to obtain summary statistics. For the population comparisons, we included an additional 1,234 mtDNA genomes from 51 Thai/Lao populations from our previous study (Kutanan et al., 2017) (Supplementary Table 1), for a total of 1,794 sequences from 73 populations (Figure 1). The matrix of genetic distances (*Φ*_*st*_, pairwise difference), Analyses of Molecular Variance (AMOVA), and a Mantel test of the correlation between genetic and geographic distances were also carried out with Arlequin. Three types of geographic distances were computed, as previously described (Kutanan et al., 2017). To get a broad picture of population relationships in Asia, we included an additional 1,936 published mtDNA genomes from 61 Asian populations (Supplementary Table 1) and calculated the *Φ* _*st*_ matrix by Arlequin.

STATISTICA 10.0 (StatSoft, Inc., USA) was used to construct a multi-dimensional scaling plot (MDS) from the *Φ* _*st*_ distance matrix. A Neighbor Joining (NJ) tree (Saitou and Nei, 1987) was also constructed from the *Φ* ^*st*^ matrix, using MEGA 7 (Kumar et al., 2016).

A Discriminant Analysis of Principal Components (DAPC) was employed using the dapc function within the adegenet R package (Jombart et al., 2011). Median-joining networks (Bandelt et al., 1999) of haplogroups without pre- and post-processing steps were constructed with Network (<u>www.fluxus-engineering.com</u>) and visualized in Network publisher 1.3.0.0.

Bayesian Skyline Plots (BSP) per population and maximum clade credibility (MCC) trees per haplogroup, based on Bayesian Markov Chain Monte Carlo (MCMC) analyses, were constructed using BEAST 1.8. BEAST input files were created with BEAUTi v1.8 (Drummond et al. 2012) after first running jModel test 2.1.7 (Darriba et al. 2012) in order to choose the most suitable model of sequence evolution. BSP calculations per population were executed with mutation rates of 1.665 ×10^-8^ (Soares et al., 2009) and Tracer 1.6 was used to generate the BSP plot from BEAST results. The BEAST runs by haplogroup were performed with the data partitioned between coding and noncoding regions with respective mutation rates of 1.708 × 10^-8^ and 9.883 × 10^-8^ (Soares et al., 2009). The Bayesian MCMC estimates (BE) and credible intervals (CI) of haplogroup coalescent times were calculated using the RSRS for rooting the tree, and the Bayesian MCC trees were assembled with TreeAnnotator and drawn with FigTree v 1.4.3.

An Approximate Bayesian Computation (ABC) approach was utilized to test different demographic scenarios concerning the relationships of SEA language families and the origin of central Thai populations. Employing an ABC methodology allowed us to simulate the evolution of complete mitochondrial sequences, by means of coalescent theory, under different competing models and to select the model that was best able to recreate the variation observed in our populations. The simulations were generated considering prior distributions associated with different model parameters. For the maternal origin of central Thai (CT) populations, we considered the same three demographic scenarios tested in our previous study for the origins of North/Northeastern Thai and Laos populations (Kutanan et al., 2017): demic diffusion; an endogenous origin (with cultural diffusion of the TK language); and admixture (Figure 2). The demic diffusion model postulates a first split of AA-speaking Mon (MO) and Khmer (KH) from the TK-speaking populations (Xishuangbanna Dai and CT) ~ 3 kya (Sun et al. 2013) followed by a later split of CT from Xishuangbanna Dai ~ 1.2 kya (O’ Connor 1995; Pittayaporn 2014) (Figure 2a). The endogenous scenario involves instead an early split of the Xishuangbanna Dai from CT and AA groups, with a later division of CT and AA ~ 0.8 kya (Baker and Phongpaichit, 2009) (Figure 2b). The admixture model incorporates the same demographic history as the demic diffusion model, but includes additional gene flow between AA and CT after the latest separation (0.8 kya) (Figure 2c). For all the models in the CT origin test, we assumed constant population sizes that were allowed to vary among groups, a fixed mutation rate (4.08 × 10^-7^) (Fu et al., 2013), and fixed separation times based on historical records, We assigned a uniform prior on the effective population size of the three groups over the interval 1,000-100,000 and on the migration rate for the admixture model between 0.01-0.2. The mtDNA genomes from CT groups (*n* = 210) were generated in the present study, while Mon (MO) sequences consisted of 49 new sequences generated in the present study plus an additional 153 MO and KH sequences reported previously (Kutanan et al., 2017). The Xishuangbanna Dai sequences were obtained from a previous study (Diroma et al., 2014)

**Figure 2.**
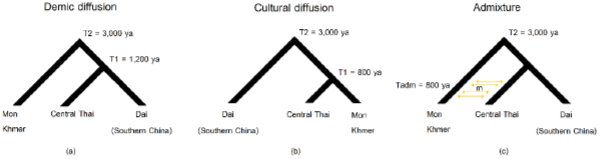
Three demographic models for the ABC analysis of CT origins: demic diffusion (a); cultural diffusion (b); and admixture (c)

For testing the genetic relationships of populations from the different SEA language families, we included populations speaking AA, AN, ST and TK languages but excluded HM because of its low population size in SEA and limited mtDNA genome data. We analyzed five tree-like demographic histories based on linguistic data for Model 1-Model 3 (Peiros, 1998; Sagart, 2004; 2005; Starosta, 2005) (Figure 3a-3c) and based on the geographic distribution of these languages for Model 4 and Model 5 (Figure 3d-3e). Since the AA, TK and ST are the languages spoken in MSEA while AN is the major language in ISEA, Model 4 and Model 5 propose a closer affinity of AA, TK and ST and set AN as an outgroup. Model 4 postulates an AA-TK affinity while Model 5 is a trifurcation of AA, TK and ST. In all the models, we assume expanding population sizes, a fixed mutation rate (4.08 × 10^-7^) (Fu et al., 2013), fixed separation times based on historical records and assigned a uniform prior distribution on both the current and ancestral effective population sizes over the range 1,000-100,000 and 1,000-50,000, respectively. We combined our Thai/Lao data with selected published mtDNA genomic data as follows: 1,219 TK sequences (present study; Diroma et al., 2014; Kutanan et al., 2017), 876 AN sequences (Gunnarsdóttir et al., 2011a; Gunnarsdóttir et al., 2011b; Jinam et al., 2012; Delfin et al., 2014; Ko et al., 2014), 627 AA sequences (present study; Kutanan et al., 2017) and 440 ST sequences (present study; Zhao et al., 2009; Zheng et al., 2011; Summerer et al., 2014; Li et al., 2015) (Supplementary Table 1). Due to the uneven sample sizes of these four groups, we simulated 440 sequences for each of the model populations as 440 sequences represents the smallest sample size; thus, the final dataset consists of 1,760 sequences.

**Figure 3.**
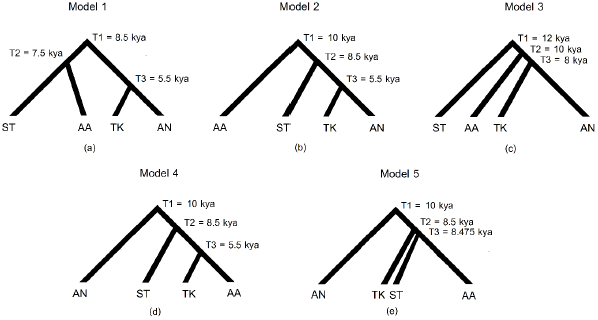
Five demographic models for the ABC analysis of the relationships of populations from four MSEA language families. Model 1 (a), Model 2 (b) and Model 3 (c) are based on Starosta (2005), Sagart (2004, 2005) and Peiros (1998), respectively, while Model 4 (e) and Model 5 (f) are based on the present geographic distributions of the languages (ISEA for AN and MSEA for ST, TK and AA); see text for further details.

Because of the computational cost of simulating a large number of complete mitochondrial sequences, we utilized a novel approach (Pudlo et al., 2016) based on a machine learning tool called “Random Forests” (Breiman, 2001). This new method can greatly reduce the number of simulations required to select the corrected model from a set of competing ones. ABC-Random Forests uses a machine-learning algorithm (based on a reference table of simulations) to predict the most suitable model at each possible value of a set of covariates (i.e. all summary statistics used to summarize the data). Random forest uses a classification algorithm which allows one to overcome the difficulties in the choice of the summary statistics, while also gaining a larger discriminative power among the competing models (see details in Pudlo et al. 2016).

To generate the simulated datasets, we used the software package ABCtoolbox (Wegmann et al. 2011) running 10,000 simulations for each model. We computed a set of summary statistics using arlsumstat (Excoffier & Lischer, 2010) describing both within-population (number of haplotypes, haplotype diversity, total and private number of segregating sites, Tajima’ s D, and average number of pairwise differences for each population), and between-population diversity (Φ _*st*_ and mean number of pairwise differences between populations). We randomly resampled 440 sequences from AA, AN and TK groups before computing the summary statistics for the observed data, so as to make them comparable with the simulated data.

## Results

### Genetic diversity and relationships

We generated 560 complete mtDNA sequences with mean coverages ranging from 54× to3687X (GenBank accession numbers will be provided upon acceptance) and identified 412 haplotypes. Genetic diversity values were lowest in the Karen group KSK2 (*h* = 0.83 ± 0.08;haplogroup diversity = 0.73 ± 0.09; *S* = 99 (Table 1)), although this was also the group with the lowest sample size. High genetic diversities were observed in CT populations (*h* = 1.00 ± 0.01 in CT2; haplogroup diversity = 0.99 ± 0.01 in CT2 and CT4; *S* = 346 in CT2) and Mon from central Thailand (MO7) (MPD = 39.32 ± 17.70 and π = 0.0024 ± 0.00119) (Table 1).

We observed 174 haplogroups among the 560 sequences; when combined with our previous study of Thai/Lao populations (Kutanan et al. 2017), there are a total of 1,794 sequences from 73 populations (Figure 1). In total there are 1,103 haplotypes and 274 haplogroups, of which 62 haplogroups were not observed in the previous study (Supplementary Table 2). An analysis of haplotype sharing (Supplementary Figure 1) shows that all four Karen groups (KSK1, KSK2, KPW and KPA) share haplotypes, indicating high gene flow among them. The Mon (MO6-MO7) shared haplotypes with several other ethnic groups, e.g. Yuan (YU) and Central Thai (CT), whereas most of the CT populations shared haplotypes more often with northeastern Thai than northern Thai groups (Supplementary Figure 1).

The AMOVA revealed that overall, 7.10% of the genetic variation is among populations (Table 2). Classifying populations by language family resulted in a slightly higher proportion of variation among groups (0.91%, *P* < 0.01) than a geographic classification (0.17%, *P* > 0.01), but for both classifications there is much more variation among populations within the same group (Table 2). Thus, neither geography nor language family is indicative of the genetic structure of Thai/Lao populations. Within each language family, the variation among AA groups (11.14%) was greater than that of ST (6.51%) or TK (4.59%) groups, indicating greater genetic heterogeneity of AA groups. Interestingly, we observed that the CT groups are the most homogenous of the TK groups, with only 1.64% of the variation among groups. However, Lue groups had higher heterogeneity (7.26%) than the average for TK groups (4.59%). A Mantel test for correlations between genetic and geographic distances indicates no correlation for all three types of geographic distances, i.e. great circle distance (*r* = 0. 0216, *P*> 0.01), resistance distance (*r* = −0.0996, *P*> 0.01) and least-cost path distance (*r* = 0.0459, *P* > 0.01), further supporting the limited impact of geography on the genetic structure of Thai/Lao populations. Furthermore, a DAPC analysis showed that clustering groups by language family resulted in more discrimination among groups than clustering by geographic criteria (Supplementary Figure 2).

**Table 2.** AMOVA results

| No. of groups | No. of groups | No. of populations | Within populations | Among populations within groups | Among groups |
| --- | --- | --- | --- | --- | --- |
| **Total<sup>b</sup>** | 1 | 73 | 92.90 | 7.10* |  |
| **AA/TK/ST<sup>b</sup>** | 3 | 73 | 92.47* | 6.62* | 0.91* |
| **Austroasiatic<sup>b</sup>** | 1 | 23 | 88.86 | 11.14* |  |
| Mon<sup>b</sup> | 1 | 7 | 93.10 | 6.90* |  |
| H'tin<sup>a</sup> | 1 | 3 | 74.29 | 25.71* |  |
| Lawa<sup>a</sup> | 1 | 3 | 92.22 | 7.78* |  |
| **Sino-Tibetan (Karen)** | 1 | 4 | 93.49 | 6.51* |  |
| **Tai-Kadai<sup>b</sup>** | 1 | 46 | 95.41 | 4.59* |  |
| Lue | 1 | 4 | 92.74 | 7.26* |  |
| Yuan | 1 | 6 | 96.10 | 3.90* |  |
| Central Thai | 1 | 7 | 98.36 | 1.64* |  |
| Khon Mueang<sup>a</sup> | 1 | 10 | 96.57 | 3.43* |  |
| Lao Isan<sup>a</sup> | 1 | 4 | 97.69 | 2.31* |  |
| Phuan<sup>a</sup> | 1 | 5 | 94.71 | 5.29* |  |
| **Geography<sup>b</sup>** | 6 | 73 | 92.85* | 6.99* | 0.17 |
| Northern<sup>b</sup> | 1 | 38 | 92.13 | 7.83* |  |
| Northeastern<sup>a</sup> | 1 | 16 | 91.29 | 8.71* |  |
| Central<sup>b</sup> | 1 | 14 | 95.84 | 4.16* |  |
| Western<sup>b</sup> | 1 | 3 | 99.12 | 0.88 |  |

\* indicates $P < 0.01$
a = data set from previous study (Kutanan et al., 2017)
b = data combined from previous and present studies to total 73 populations

The MDS showed that the most differentiated groups were two H’ tin gropus (TN2 and TN1) and Seak (SK), as found previously (Kutanan et al., 2017) and the central cloud of the plot is difficult to see population clustering trends (Supplementary Figure 3). After omitting these outlier groups, a 3-dimensional MDS provides an acceptable fit (Figure 4a-c) and shows some clustering of populations by language family (with considerable overlap). In the NJ tree (Supplementary Figure 4), Karen (KSK1 and KSK2) groups showed distant affinities with H’ tin groups (TN1-TN3), even though they reside quite far apart from one another in Northern Thailand, with multiple intervening mountain ranges. The MDS plot of Asian populations indicated that SEA groups are separated from Indian groups; some Mon groups (MO1, MO5 and MO6) are closely related to the Indian groups as well as Myanmar (BR1 and BR2) and Cambodia (KH_C and AA_C), while the other Mon (MO2-MO4, MO7) are close to the other SEA populations (Supplementary Figure 5).

**Figure 4.**
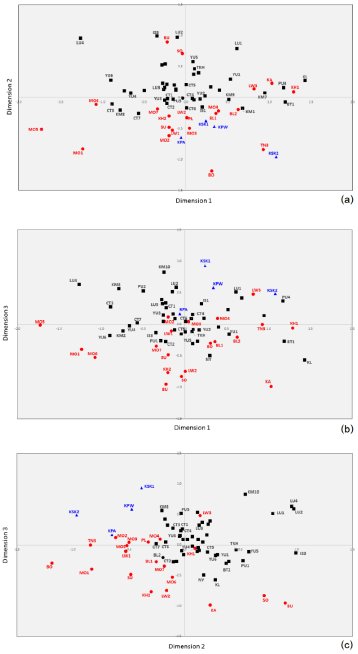
MDS plots based on the *Φ* _*st*_ distance matrix for 70 populations (after removal of three outliers: TN1, TN2, and SK). The stress value is 0.0804. Population abbreviations are shown in Supplementary Table 1.

**Figure 5.**
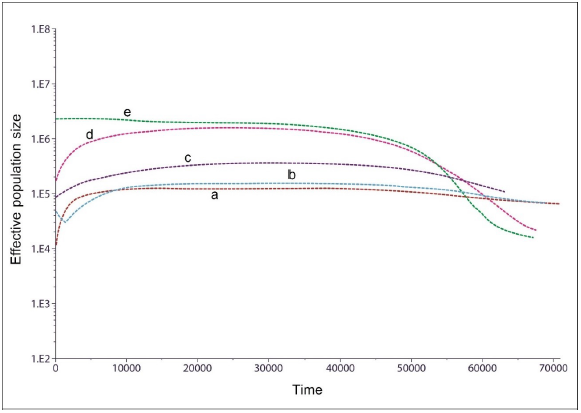
The BSP plots for 5 different trends found in 22 populations; KSK2, MO6, LU4 (a), KSK1 (b), KPA (c), KPW, MO7, KPW, TKH, LU1-LU2, YU3-YU6, CT6-CT7 (d) and LU3,CT1-CT5 (e). Population abbreviations are in Supplementary Table 1. Each line is the median estimated maternal effective population size (y-axis) through time from the present in years (x-axis).

### MtDNA haplogroups

Fourteen of the 174 haplogroups occur in at least ten individuals and together account for 33.92% of the 560 sequences; these are F1a1a, B6a1a, F1f, B5a1a, F1a1a1, C7a1, C7a, M^*^, M12a1a, M21a, M7b1a1a3, R9b1a1a, R9b1a3 and B5a1b1 (Supplementary Table 2). These common haplogroups are mostly prevalent in AA groups (e.g. M^*^ and M12a1a in MO6, 50.00%)and ST-speaking Karen groups (B6a1a, C7a1, R9b1a1a in KSK1, 84.00%; F1a1a in KSK2, 46.15%; F1a1a, C7a1, R9b1a1a in KPW, 70.83%; B6a1a, F1a1a1, M^*^ and M21a in KPA,56.00%). These very distinct haplogroup distributions further emphasize the genetic distinctiveness of AA and ST groups.

The remaining haplogroups (66.08%), which occur in lower frequency, tend to be more widely distributed, e.g. G2a1 and basal M sublineages in MO7 and subhaplogroups F (x F1a1a and F1a1a1), M7b1a1 and B4 in Lue (LU) and Khuen (TKH) at varying frequencies, (Supplementary Table 2). New subhaplogroups of B4 (B4a1a, B4a1c2, B4b1c1, B4c, B4c2c, B4g2 and B4m), F3 (F3a, F3b, F3b+152) and M7 (M7b1a1g, M7b1a1h, M7c1c3 and M7c2b) are present mostly in TK populations (Supplementary Table 2). In agreement with the AMOVA results (Table 2), the CT groups were more similar in haplogroup distribution. The CT groups show a wide haplogroup distribution with various haplogroups occurring in a few individuals and very few haplogroups at high frequency (most are lower than 10%). Several subclades of M lineages (M12a2, M12b2, M13b1, M17c1a1, M17c1a1a, M21b2, M2a1a, M32’ 56, M37e2, M50a1,M51a1a, M73a1, M73b, M7, M7b, M7b1a1g, M7c1c3, and M7c2b) are newly-reported in Thai/Lao groups and are exclusively found in CT populations. Interestingly, other new haplogroups, e.g. R11’ B6, R21, R23, U1a1c1a, U1a1c1d, U2a1b and U2a2 were also observed in the CT groups (Supplementary Table 2).

In the combined Thai/Lao dataset, SEA specific haplogroups (B, F and M7) are prevalent in almost all groups (overall frequency 55.18%), with the exception of some AA groups (i.e. Mon, Suay, Nyahkur, Khmer and Lawa), Karen, and CT groups; these groups have other widespread haplogroups, e.g. D, M12-G, M (xM12-G, M7), A, C and N (xN9a) (Figure 1). Networks of common SEA specific haplogroups, e.g. B5a, F1a, F1f and M7b, tend to exhibit star-like structures, indicative of population expansions (Supplementary Figure 6). Apart from F1a1a (xF1a1a1), other more-prevalent haplogroups of Karen (B6a1a and C7a1) do not show indications of population expansion, but rather sharing of sequences, suggesting population contraction (Supplementary Figure 6). Apart from B and F1, other lineages, that is, C7a1 and A17 and N8 which are sublineages of C, A and N (xN9a), respectively are observed in the Karen (Figure 1). Haplogroup C7 had a very high frequency in northeast Asia and eastern India (Derenko et al., 2010) while haplogroup A was previously reported to be specific to North and Central Asia (Derenko et al. 2007). A high proportion of C and A lineages were previously observed in ST-speaking Barmar and Karen from Myanmar (Summerer et al., 2014). For the TK-specific haplogroups, i.e. B4 and M7c, there was no obvious signal of population expansion in the networks (Supplementary Figure 6).

For the combined dataset, we estimated coalescence ages of SEA haplogroups and their sublineages. We analyzed haplogoups that have additional sequences from the present study and have more than five sequences in total (Table 3). The ages of major haplogroups are generally consistent with previous studies (Kutanan et al., 2017). However, we obtained more data from several sublineages which were not dated previously, e.g. B4c1b, B6a1, C4, C7a, D4a, F1c, F1e, F1g, F2, F3, F4a2 and G2a (Table 3).

**Table 3.**
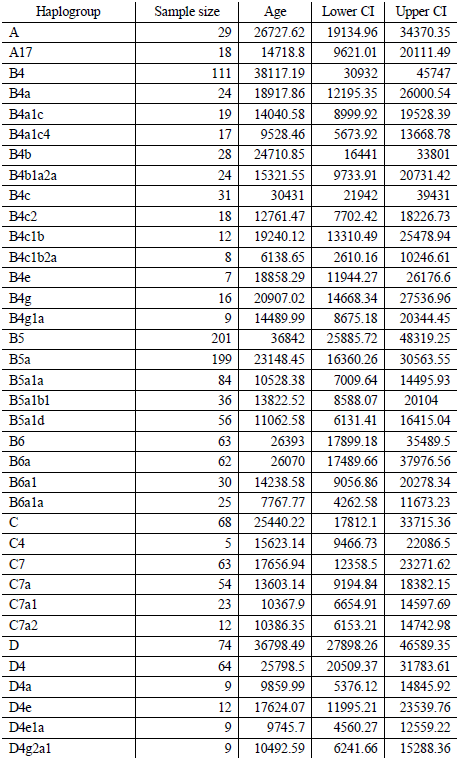
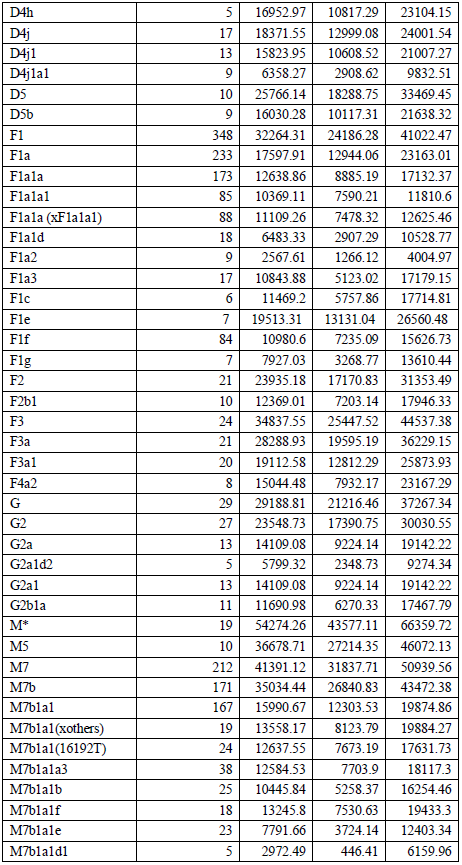
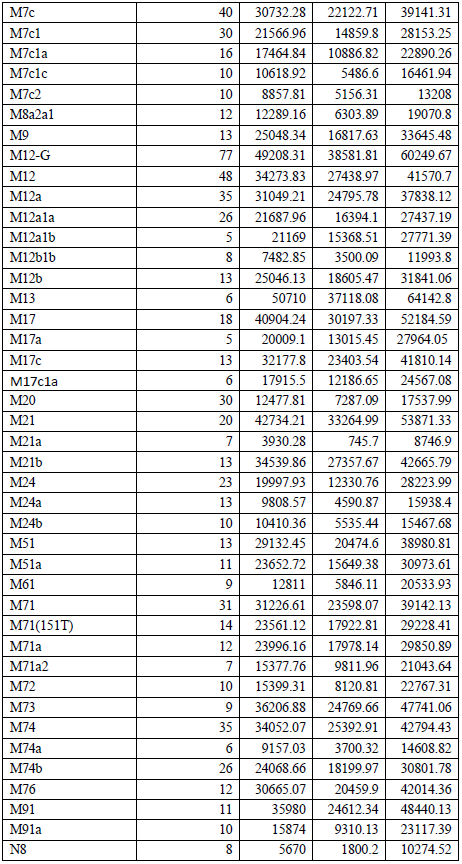
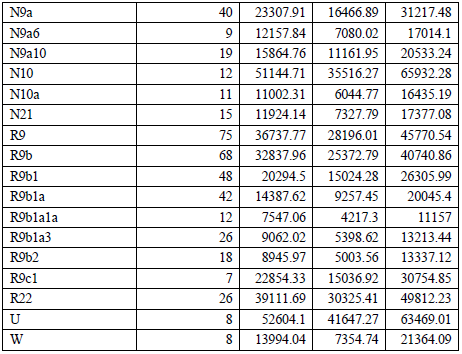
Coalescent ages based on Bayesian estimation with 95% credible interval (CI) and using the 1,794 Thai/Lao mtDNA sequences.

There are many archaic lineages with ages older than 30 kya that found in our Thai/Lao samples, e.g. B4, B5, D, F1, F3, M7, M^*^, M12, M13, M17, M21, M71, M73, M74, M91, R9, R22,N10 and U. Many of them are major lineages and distributed in our Thai/Lao samples as well as in other SEA populations, and have been previously discussed (see details in Kutanan et al., 2017). Here, we focused on some uncommon ancient lineages, i.e. M^*^, M17, M21, M71, M73, M91 and U. Nineteen sequences were classified as superhaplogroup M^*^ (i.e., they could not be classified into other M sublineages) and date to ~ 54.27 kya; most of them occur in the Mon (52.63%) and Karen (KPA) (15.79%). M17 bifurcated to M17a and M17c ~ 40.90 kya, which 61.11% is contributed by the Central Thai (CT). M17a is proposed to be an early mtDNA lineage, which putatively originated in MSEA and migrated to ISEA (Belwood, 2017; Tumonggor et al., 2013) while M17c was found in the Philippine populations (Tabbada et al., 2010; Delfin et al., 2014). We here date these lineages to ~ 29.02 kya (M17a) and ~ 32.18 kya (M17c) (Table 3). M21 bifurcates ~ 42.73 kya to the older clade (M21b) and younger clade (M21a) with ages 34.54 kya and 3.93 kya, respectively. M21b was found in AA-speaking and CT groups whereas M21a is new lineage in Thai/Lao populations, found in the Karen and MO7. M21a is most common among the Semang and M21b is found in both the Semang and Senoi from Malaysia (Hill et al., 2006). Two major sublineages of M71 are M71(151T) and M71a. Although M71 is rare (~ 0.02%) in our study, its frequency is higher than reported previously in MSEA (Peng et al., 2010; Bodner et al., 2011; Zhang et al., 2013) and ISEA (Tabbada et al., 2010). The estimated divergence time of M71 ~ 31.22 kya, slightly lower than a previous estimate of ~ 39.40 kya (Peng et al., 2010). The ages of M71(151T) and M71a are ~ 23.56 kya and ~ 24.00 kya. About 50% of M71a is from CT individuals, with the remainder found in other TK groups and in the Blang, an AA group. M73 was mostly contributed by the MO (44.44%) and CT (44.44%). It was also reported previously at low frequency in MSEA (Peng et al., 2010; Bodner et al., 2011; Zhang et al., 2013) and ISEA (Tabbada et al., 2010). We dated this lineage to ~ 36.21 kya, consistent with a previous estimate of ~ 37.80 kya (Peng et al., 2010). Notably, M17, M21, M71 and M73 are ancient maternal lineages of SEA found in both MSEA and ISEA, reflecting linkages between the early lineages in SEA (Jinam et al., 2010).

M91, dated to ~ 35.98 kya, is another proposed indigenous SEA haplogroup. The age estimated here is slightly lower than in a previous study of Myanmar (~ 39.55 kya) (Li et al., 2015). A sublineage, M91a, dates to ~ 15.87 kya and is found in MO, Karen (KPA) and CT (Supplementary Table 2). Interestingly, haplogroup U is the second oldest lineage in this study with an age of ~ 52.60 kya, which is slightly higher than a recent estimate of 49.60 kya (Larruga et al., 2017). Subhaplogroups U1 and U2, which are restricted to CT groups, are autochthonous to the Near East (Derenko et al., 2013) and South Asia (Palanichamy et al., 2004), respectively. Overall, the CT groups contrast with other Thai/Lao groups in exhibiting several ancient haplogroups (especially basal M lineages) at low frequency.

Finally, several haplogroups associated with the Austronesian expansion from Taiwan, namely B4a1a1a, M7b3, M7c3c, E1a1a and Y2 (Peng et al., 2010; Duggan et al., 2014; Ko et al., 2014; Soares et al., 2016) were not observed, further supporting that this expansion had at most a limited impact on mtDNA lineages in MSEA.

### Bayesian skyline plots

Bayesian skyline plots (BSP) of population size change over time were constructed for each group, and five typical patterns were observed (Figure 5). The four Karen populations all showed different patterns: KSK2 (and also MO6 and LU4) displayed unchanged population size until ~ 1–2 kya followed by sharp reductions (Figure 5, pattern a); KSK1 was also constant in size, with a sudden increase in the last 1-2 kya (Figure 5, pattern b); KPA was basically constant in size over time (Figure 5, pattern c); and KPW exhibited the most common pattern (also observed in MO7, KPW, TKH, LU1-LU2, YU3-YU6, CT6-CT7), consisting of population expansion between 50-60 kya, followed by a decrease in the last 5 kya (Figure 5, pattern d). Finally, population growth without further change was found for LU3 and CT1-CT5 (Figure 5, pattern e). The BSP plots for each individual population are depicted in Supplementary Figure 7.

### Demographic models for the origin of central Thai people

In our previous study we used demographic modeling to show that northern and northeastern Thai groups most likely originated via demic diffusion from southern China (Kutanan et al. 2017). Here we use the same approach to test three demographic scenarios concerning the origins of central Thai groups: (1) descent from the prehistorical Tai stock of southern China via demic diffusion, like their neighbors in northern and northeastern Thai (Figure 2a); (2) local AA groups (Mon and Khmer) who changed their identity and language via cultural diffusion to become TK groups (Figure 2b); or (3) descent from a migration from southern China that admixed with the local Mon and Khmer people (Figure 2c). The LDA plot shows that the observed data fall within the distribution of simulated data under the three models, indicating a plausible result for the simulated data (Supplementary Figure 8). The demic diffusion model had the highest posterior probability at 0.604 and also selected slightly more often among the classification trees (0.515) than the admixture model (0.404); both of them were selected much more often than the model of cultural diffusion (0.081). We conclude that demic diffusion, possibly with some admixture, is the most likely scenario for the origins of central Thai populations.

### Genetic relationships of populations from different language families

We also used the demographic modeling approach to test different models for the genetic relationships of populations belonging to the four main SEA language familes (TK, AA, AN and ST). In doing so, it is important to keep in mind that we are not testing the relationships of these language families, as that would require linguistic data. However, determining the best-fitting model based on genetic relationships may help discriminate among hypotheses concerning the language family relationships that make predictions about the genetic relationships of populations speaking those languages. We tested five models of the language family relationships (Figure 3). The observed data fall within the range of the simulated data in the LDA plot (Supplementary Figure 8). The model that best fit the mtDNA genome data was Model 1, according to Starosta (2005) (Figure 3a). The posterior probability of this model is 0.657, and it was selected slightly more often among the classification trees (0.509) than Model 2 (0.311); the other models were much less often selected among the classification trees (0.037 for Model 3; 0.112 for Model 4;0.031 for Model 5). Because of high selection frequency of Model 1 and Model 2, which have in common can ancestral relationship of TK and AN groups (Figure 3a and 3b), we conclude that the TK and AN groups are descended from a common ancestral population.

## Discussion

The present study adds to our previous study of Thai/Lao mtDNA genome sequences by including 22 additional groups from Thailand, including the AA-speaking Mon (MO), ST-speaking Karen, and several TK speaking groups, especially the central Thais (CT). The Mon who were a previously dominant group in MSEA with centers in the present-day southern Myanmar and central Thailand since the 6^th^ to 7^th^ century AD (Saraya, 1999), have been reported to link with Indian populations with some haplogroups, i.e. W3a1b (Kutanan et al., 2017). With data from an additional two Mon groups, there is still support for a connection between India and the Mon in the distribution of M subhaplogroups characteristic of South Asia or the Near East, e.g. M6a1a, M30, M40a1, M45a and I1b (Chandrasekar et al., 2009; Olivieri et al., 2013; Silva et al., 2017) (Supplementary Table 2). Genetic relationship analysis also reveals some Mon populations (MO1, MO5, MO7) clustering with Indian groups (Supplementary Figure 5). Thus, based on the many older mtDNA lineages observed, the modern Mon from both Thailand and Myanmar could be one of the important groups for further studies to reconstruct early SEA genetic history.

The Karen in Thailand are refugees who migrated from Myanmar starting from the 18^th^ century A.D. due to the influence from Burmese (Grundy-Warr *et al*., 2003). However, the ancestors of the Karen probably migrated from some unknown location to Myanmar, as the Karen languages are thought to have originated somewhere in north Asia or in the Yellow River valley in China, i.e. the homeland of ST languages (LaPolla, 2001). In agreement with previous studies of either different Karen subgroups or different genetic markers (Kutanan et al., 2014; Listman et al., 2011; Summerer et al., 2014), we find both northeast and southeast Asian components in the maternal ancestry of the Karen.

The present results emphasize the common maternal ancestry of central Thais (CT) and other TK speaking groups in MSEA, e.g. Laos and Southern China. Demic diffusion is still the most probable scenario for TK-speaking populations (Figure 2a), possibly accompanied by some admixture with autochthonous AA-speaking groups. It seems that the prehistoric TK groups migrated from the homeland in south/southeast China to the area of present-day Thailand and Laos, and then split to occupy different regions of Thailand, expanding and developing their own history. During the migration and settlement period, genetic intermingling with the local AA people was certainly limited, but nonetheless the modeling results, haplogroup profiles and genetic diversity values all suggest some degree of admixture in the CT groups (Supplementary Table 2; Table 1); Y chromosome and genome-wide data could provide further evidence for admixture. However, in sum, cultural diffusion did not play a major role in the spread of TK languages in SEA.

Finally, we used simulations to test hypotheses concerning the genetic relationships of groups belonging to different language families. We found that Starosta’ s model (Starosta, 2005) provided the best fit to the mtDNA data; however, Sagart’ s model (Sagart, 2004; 2005) was also highly supported. These two models both postulate a close linguistic affinity between TK and AN. Although genetic relatedness between TK and AN groups has been previously studied (Dancause et al., 2009; Mirabal et al., 2013; Kutanan et al., 2017), this is the first study to use simulations to select the best-fitting model. Our results support the genetic relatedness of TK and AN groups, which might reflect a postulated shared ancestry among the proto-Austronesian populations of coastal East Asia (Bellwood, 2006).

Specifically, the model suggests that after separation of the prehistoric TK from AN stocks around 5-6 kya in Southeast China, the TK spread southward throughout MSEA around 1-2 kya by demic diffusion process with increment of their population size without (or with possibly minor) admixture with the autochthonous AA groups. Meanwhile, the prehistorical AN ancestors entered Taiwan and dispersed southward throughout ISEA, with these two expansions later meeting in western ISEA. The lack of mtDNA haplogroups associated with the expansion out of Taiwan in our Thai/Lao samples has two possible explanations: either the Out of Taiwan expansion did not reach MSEA (at least, in the area of present-day Thailand and Laos); or, if the prehistoric AN migrated through this area, their mtDNA lineages do not survive in modern Thai/Lao populations – thus ancient DNA studies in MSEA would further clarify this issue.

## Acknowledgements

We would like to acknowledge participants and coordinators for assistance in collecting samples. We thank Prof. Murray Cox for his discussion and suggestions. This study was supported by the Max Planck Institute for Evolutionary Anthropology, the Thailand Research Fund (Grant No. MRG5980146) and Research and Academic Affairs Promotion Fund (RAAPF) of Faculty of Science, Khon Kaen University.

## Author Contribution

WK and MS designed the study. WK, JK, MSr, SR, DK and PP were involved in sample recruitment. WK, RS AH, EM, and LA generated sequencing data. WK, SG, and AB analysed the data. WK, SG, AB AH, EM, LA and MS wrote the manuscript with input from all other authors.

